# Synergy between HA cleavage site sequence and NA-mediated plasminogen recruitment as a virulence mechanism for low pathogenic avian influenza

**DOI:** 10.1101/2025.07.11.664276

**Authors:** Hui Min Lee, Kate Sutton, William Harvey, Rute Maria Pinto, Eleanor Gaunt, Samantha Lycett, Sjaak de Wit, Lonneke Vervelde, Paul Digard

## Abstract

An outbreak of H3N1 low pathogenic avian influenza virus (LPAIV) in Belgium in 2019 caused unexpected levels of mortality and morbidity in poultry. These viruses possess an NA polymorphism associated with plasminogen binding, as well as an unusual sequence around the HA cleavage site; accordingly, HA cleavage mediated by NA-driven plasminogen recruitment has been proposed to underly their systemic spread and pathogenicity. To test this, we established a reverse genetics system for A/chicken/Belgium/460/2019 and created single mutations in HA (K345R), and NA (S122N) that restored the viruses to normal consensus, as well as an HA/NA double mutant. Confirming previous work, trypsin-independent spread and HA cleavage of wild type Ck/Belgium was observed in the presence of fetal bovine serum containing plasminogen *in vitro*. Dose-dependent HA cleavage and trypsin-independent spread was also observed in the presence of purified chicken plasminogen. Compared to wild type virus, both HA cleavage and virus spread *in vitro* were reduced by the HA K345R mutation and further blocked by NA mutation S122N. Plasminogen-mediated HA cleavage was seen in a variety of avian cell lines and chicken organoids, excluding cell type-dependent effects. Furthermore, *in ovo* tests showed that mutant viruses unable to recruit plasminogen were less able to replicate systemically in chicken embryos. Bioinformatics analyses revealed other viruses which could potentially recruit plasminogen, including two independent outbreaks of H6N1 viruses, one of which we confirmed PLG-driven spread *in vitro*. We conclude that PLG-recruitment by NA is a general virulence mechanism of N1 LPAIVs

**Author summary:** Avian influenza viruses (AIV) are divided into two broad categories – high or low pathogenicity – based on the sequence of their haemagglutinin (HA) and their lethality in chickens. The majority of AIV strains circulating in the wild are low pathogenicity both in waterfowl and when they spill over into domestic poultry. However, some low pathogenicity strains can cause severe disease in poultry despite not being classified as H5 or H7 high pathogenic AIVs with an HA polybasic protease cleavage site. A severe 2019 outbreak of an H3N1 strain has been suggested to result from the neuraminidase (NA) of the virus recruiting cellular plasminogen to proteolytically activate HA. Here, we tested this hypothesis by using reverse genetics to mutate the virus in a way predicted to block this. We found that indeed, the sequence of the NA at position 122 is the primary determinant of plasminogen-driven HA cleavage but that the unusual sequence at the HA cleavage site of the outbreak virus also contributes to pathogenicity. Furthermore, we show that N1 NA sequence can be used to identify other unexpectedly virulent strains of AIV. This work therefore adds to our ability to risk assess AIV strains from sequence-based surveillance.

## Introduction

Avian influenza viruses (AIV) are classified into high pathogenic (HPAI) or low pathogenic (LPAI) viruses based on their hemagglutinin (HA) subtype and sequence near the HA cleavage site. Almost without exception, HPAI viruses (HPAIVs) are H5 or H7 subtypes that possess a polybasic cleavage site (PBCS), whereas LPAI viruses are all other subtypes and H5 or H7 subtypes without a PBCS (1, 2). In gallinaceous poultry such as chickens and turkeys, HPAIVs show systemic spread and rapidly fatal disease, whereas infection by LPAI viruses is largely restricted to the respiratory tract and intestinal organs and often causes few or no clinical signs (3). However, not all LPAI viruses are phenotypically low pathogenic; certain strains can be highly virulent, spread systemically in the animals and lead to an overt outbreak (4–10).

In 2019, an LPAI H3N1 outbreak involving various poultry species started in Belgium and spread regionally for four months, including to three farms in France (9). The outbreak was notable for its association with dramatic drops in egg production, tropism for the reproductive tract and high mortality in chickens (9). Our laboratory study using an H3N1 isolate from the outbreak, A/chicken/Belgium/460/2019 (Ck/Belgium), showed 58% mortality and 100% drop in egg production in 34-week-old specific pathogen free (SPF) laying hens (8). Virus was also detected in the oviducts of the hens at 9 to 10-days post-infection (8). This indicated that the H3N1 strain was indeed a virulent LPAIV capable of systemic spread in poultry. Therefore, it was of interest to identify the molecular reasons that contribute to the increased virulence of Ck/Belgium, both to understand the reasons for it and to enhance risk prediction of future novel AIV strains as they arise.

The surface of an influenza virus particle contains two glycoproteins: HA and neuraminidase (NA). In order to activate virus infectivity and permit cell entry, the HA precursor polypeptide (HA0) must be cleaved by host proteases into two functional subunits, HA1 and HA2 (11–13). Without this specific proteolysis event freeing the N-terminus of the HA fusion peptide, viral and host membranes cannot be fused and virus entry fails. Therefore, HA sequence around the HA cleavage site can be an important determinant of virus pathogenicity in different hosts (2, 14). The PBCS in HPAIV HAs are cleaved intracellularly by ubiquitously expressed furin family proteases (15, 16), whereas the HAs from LPAI viruses with a monobasic cleavage site are usually activated extracellularly by secreted trypsin-like proteases found in the airway or gut lumens, but not within the internal organs (12, 17). An additional mechanism for HA cleavage which depends on the sequence of the viral NA has been found in certain strains of influenza A virus. In a human-isolated but mouse-adapted H1N1 strain, A/Wilson Smith Neurotropic/1933 (WSN), the presence of a C-terminal lysine and loss of glycosylation site at position 130 (WSN numbering) on NA leads to NA binding to plasminogen (PLG), which is subsequently converted to plasmin that cleaves adjacent HA molecules (18, 19). PLG recruitment by NA in WSN has been associated with increased pathogenicity in mice, including neurotropism, presumably because of the systemic availability of PLG (18). Consistent with this, PLG-knockout mice infected with another laboratory-adapted H1N1 strain, A/Puerto Rico/8/1934 (PR8), which is also able to use PLG for HA cleavage, showed higher survival and lower weight loss compared to wild type animals (20).

Notably, the NA from the H3N1 Ck/Belgium outbreak also lacks the equivalent glycosylation site (at position 122) to the WSN NA and a previous study has shown that another isolate from the outbreak, A/chicken/Belgium/1940/2019, exhibited trypsin-independent spread in cell culture in the presence of PLG-containing serum (21). Furthermore, virus replication was blocked by a plasmin inhibitor, 6-aminohexanoic acid (6-AHA), suggesting a mechanism for the increased virulence of the Ck/Belgium family viruses (21). In addition, the Ck/Belgium HA also contains an unusual monobasic cleavage sequence, with a lysine (K) at the HA1 boundary (P1 position), instead of the more common arginine (R) residue seen in H3 HAs (Fig 1A). A previous study using WSN has reported that a non-consensus amino acid at the P2 position of the HA cleavage site affected virus virulence under plasminogen-dependent conditions (22), indicating the possibility of collaboration between the atypical HA cleavage site and PLG binding by NA (21, 22). Accordingly, we set out to test the molecular hypotheses that the unexpectedly high virulence of Ck/Belgium lineage viruses resulted from plasminogen recruitment by NA working in concert with a modified HA monobasic cleavage site. We confirmed PLG-mediated cleavage of the Ck/Belgium HA in the context of virus infection and found that the atypical NA and HA sequences combine for efficient PLG-mediated spread in a variety of avian cell systems. We also identified PLG-mediated spread in a separate H6N1 outbreak, suggesting that this may be a general pathogenicity mechanism for LPAIV virulence.

**Figure 1.**
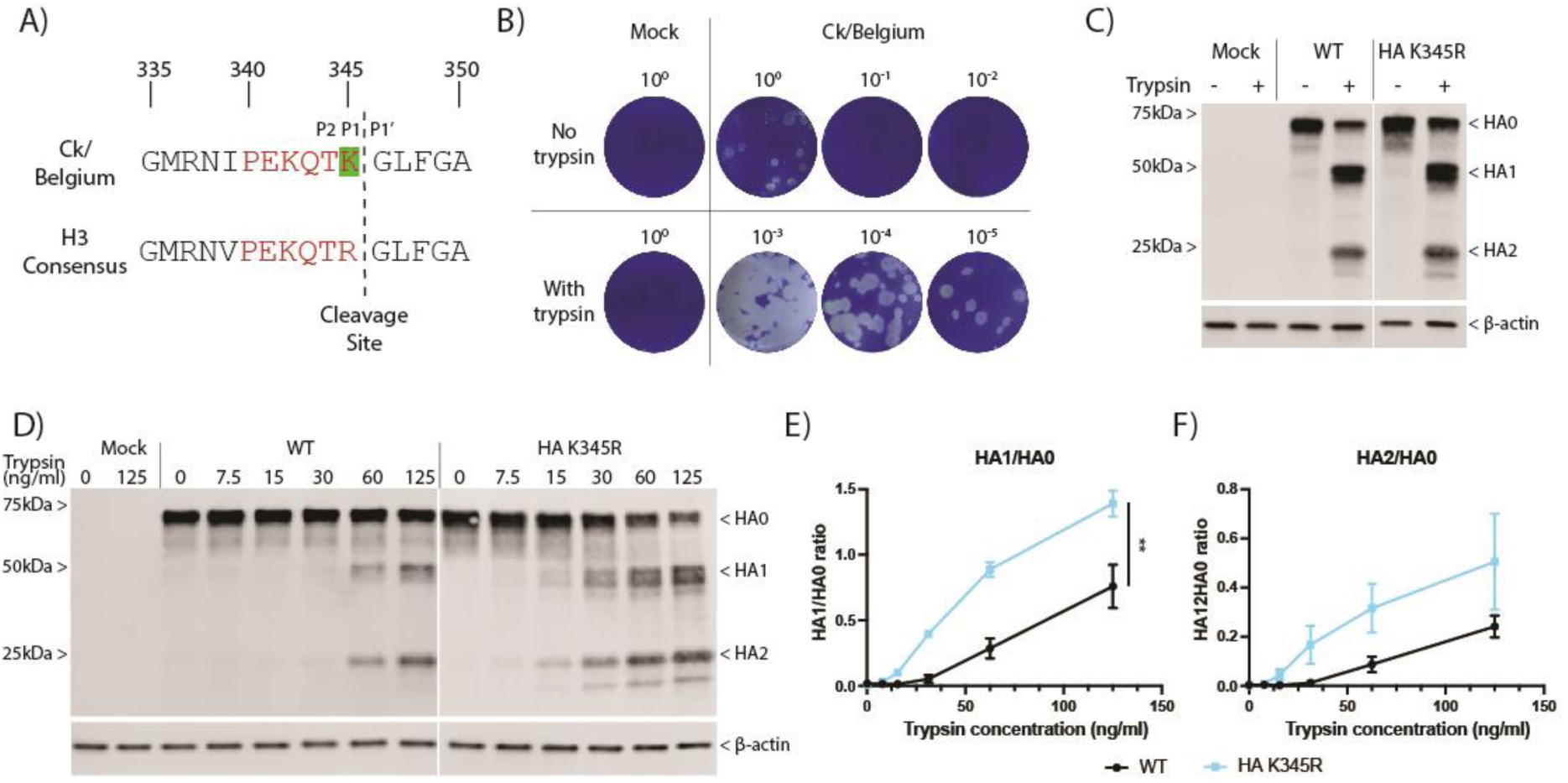
Trypsin-dependency of CK/Belgium in cell culture. (A) Alignment of HA sequences near the cleavage site for Ck/Belgium and an H3 consensus. Red lettering indicates the cleavage site motif while the amino acid highlighted in green at position 345 indicates the polymorphism of Ck/Belgium in this motif compared to the H3 consensus. (B) Plaque titration in MDCK cells (with trypsin) of supernatant collected from mock or Ck/Belgium-infected CLEC213 cells (MOI=0.001) incubated in the presence or absence of trypsin for 48 hours. (C) CLEC213 cells were infected with WT or HA K345R at MOI of 3 for 16 hours and 1 μg/ml trypsin was added for the last 2 hours of infection. Cell lysates were examined by SDS-PAGE and western blotting for HA and ß-actin. Positions of molecular mass markers are indicated on the left. (D) CLEC213 cells were infected with WT or HA K345R viruses at MOI of 3 for 16 hours and the indicated trypsin concentrations were added for the last 2 hours of infection. SDS-PAGE and western blotting were performed to detect HA and ß-actin. Positions of molecular mass markers are shown on the left. (E, F) Ratios of cleaved (HA1 or HA2) to uncleaved HA (HA0) products were quantified by densitometry and data are pooled data from 2 each independent infection and transfection experiments expressed as mean ± SEM. Unpaired t-test of area under the curve for each condition was used for statistical analyses. ** = p < 0.01.

## Materials and Methods

### Cells

Chicken ANP32A-expressing Madin-Darby canine kidney cells (MDCK/ANP32A; a kind gift from Prof Massimo Palmarini, University of Glasgow Centre for Virus Research, UK), quail fibroblast cells (QT-35; a kind gift from Dr Laurence Tiley, University of Cambridge, UK) (23) and human embryonic kidney cells (HEK293T; ATCC) were cultured in Dulbecco’s modified Eagle’s medium (DMEM; Sigma) supplemented with 10% fetal bovine serum (FBS; Gibco), 100 U/ml penicillin and 100 µg/ml streptomycin (Gibco). 1 µg/ml puromycin was added for the selection marker in MDCK/ANP32A cells. Chicken lung epithelial cells (CLEC213; a kind gift from Dr Sascha Trapp, French National Institute for Agriculture, Food and Environment, France) (24) were cultured in DMEM-F12 (Sigma) supplemented with 8% FBS, 100 U/ml penicillin and 100 µg/ml streptomycin. Chicken embryonic fibroblast cells (DF-1) (25) and duck embryonic fibroblast cells (CCL-141; a kind gift from Dr Leah Goulding, University of Nottingham, UK) were cultured in DMEM-F12 supplemented with 10% FBS, 100 U/ml penicillin and 100 µg/ml streptomycin.

### Antibodies and antisera

The following primary antibodies were used: rabbit anti-H3 for A/HKx31 (H3N2; a kind gift from Prof Janet Daly, University of Nottingham, UK), laboratory-made rabbit anti-nucleoprotein (NP; 2915) (26), goat anti-influenza A H1N1 (5315-0064,Bio-Rad), rat anti-α-tubulin (clone YL1/2, Invitrogen) and mouse anti-ß-actin (ab8226, Abcam). Secondary antibodies used were: Alexa Fluor 800-conjugated goat anti-rabbit IgG (Invitrogen), Alexa Fluor 680-conjugated goat anti-rat IgG (Invitrogen), IRDye 594RD goat anti-rabbit IgG (LI-COR Biosciences) and IRDye 680RD donkey anti-mouse IgG (LI-COR Biosciences).

### Viruses and reverse genetics

H3N1 A/chicken/Belgium/460/2019 (Ck/Belgium) (8) and H3N2 A/Udorn/307/1972 (Udorn; a kind gift of Dr Richard Compans, Emory University, Atlanta) (27) have been previously described. H1N1 A/Wilson Smith Neurotropic/1933 (WSN) and H6N1 A/chicken/Netherlands/917/2010 were sourced from the Division of Virology, Department of Pathology, University of Cambridge and Royal GD virus strain collections respectively.

To establish a reverse genetics system for Ck/Belgium, individual full-length segments corresponding to an in-house sequence of the coding regions of A/chicken/Belgium/460/2019 clone GD4 (hereafter Ck/Belgium) with the missing untranslated regions obtained by deriving an H3N1 consensus sequence were ordered from Genewiz, UK and subcloned into a bi-directional RNA polymerase I/II “pDUAL” reverse genetics vector (28) at the *BsmBI* restriction site. Each segment was sequenced to confirm identity. Desired mutations were introduced using a QuikChange Lightning Site-Directed Mutagenesis kit (Agilent) according to manufacturer’s instructions. Wild type Ck/Belgium and its mutant derivatives were then produced by reverse genetics as previously described (29). Briefly, HEK293T cells were transfected with eight pDUAL plasmids encoding each segment of influenza virus. Following overnight incubation, medium was replaced with serum-free DMEM supplemented with 0.14% bovine serum albumin (BSA; Merck) and 1 μg/ml L-(tosylamido-2-phenyl) ethyl chloromethyl ketone (TPCK)-treated trypsin (Worthington Biochemicals), and co-cultured with MDCK/ANP32A cells to obtain efficient rescue. Cell culture supernatant was collected at 3 days post-transfection and clarified before being used to infect 10-day old embryonated chicken eggs. Eggs were chilled overnight at 2 days post-infection, allantoic fluid harvested, clarified and titrated by plaque assays on MDCK/ANP32A cells. The presence or absence of specific mutations, as well as the receptor binding site of the HA were confirmed by sequencing.

All reverse genetics work used a loss-of-function approach and was carried out under a license (GMRA1811) from the UK Health & Safety Executive. All work with avian viruses was carried out in a segregated BSL2 laboratory in the Roslin Institute, separate from where mammalian strain of influenza A virus are handled.

### Plaque assay

Confluent cell monolayers of MDCK/ANP32A cells were used to measure virus titre. Cells were infected with ten-fold serial dilutions of virus for an hour at 37°C and then overlaid with serum-free DMEM containing 0.14% BSA, 1 μg/ml TPCK-treated trypsin and 0.8% Avicel (Dupont). Following 3 days incubation at 37°C, cells were fixed with neutral-buffered formalin for 20 minutes and stained with 0.1% toluidine blue (Sigma) for another 20 minutes.

### Immunofluorescence

Cells were infected with viruses at the MOIs stated in the figure legends and incubated in DMEM-F12 in the absence or presence of 10% FBS for 24 or 72 hours. Cells were then fixed with phosphate-buffered saline (PBS)/4% formaldehyde (Thermo Fisher Scientific) for 20 minutes, washed 3 times with PBS, blocked for 30 minutes with 1% FBS in PBS and incubated with primary antibody for an hour. Following 3 PBS/1% FBS washes, cells were incubated with species-specific secondary antibody for an hour and subsequently Hoechst dye (to stain nuclei) incubation for 10 minutes. Cells were washed 3 times with PBS before mounting with ProLong Gold antifade reagent (Invitrogen) and visualised using a DMRB fluorescence microscope (Leica).

### Immunoblotting

Cells were infected with viruses or transfected with plasmids and then incubated in the absence or presence of 10% FBS, 6-AHA (Sigma) or chicken PLG (Abcam) according to experimental design. Cells were lysed in 2x homemade Laemmli buffer (30) and separated by 4-20% SDS-PAGE (Bio-Rad). Proteins were transferred onto nitrocellulose membranes (Thermo Fisher Scientific) using a Transblot Turbo semi-dry transfer system (Bio-Rad).

Membranes were blocked with Intercept blocking buffer (Li-COR Biosciences) for 1 hour before overnight incubation with primary antibodies at 4°C. Following 3 washes in PBS/ 0.1% Tween 20, membranes were incubated with species-specific secondary antibodies for an hour. Membranes were washed 3 times and visualised using an Odyssey Fc imaging system (Li-COR Biosciences). Densitometric analyses were performed using Image Studio Lite Software (Li-COR Biosciences).

### Flow cytometry

Cells were infected with viruses at the MOIs stated in the figure legends and incubated in the absence or presence of chicken PLG according to experimental design. Cells were fixed with PBS/4% paraformaldehyde for 20 minutes, washed 3 times in PBS and incubated with primary antibody diluted in cell staining buffer (Biolegend) for 1 hour. Cells were washed 3 times in cell staining buffer and incubated with species-specific secondary antibody for 1 hour. After 3 PBS washes, the percentage of virus-infected cells was determined using a LSR Fortessa X-20 flow cytometer (BD Biosciences). 10,000 events were recorded by gating on live single cells. Data analysis was performed using FlowJo (BD Biosciences).

### Chicken embryo pathogenesis model

Pathogenesis of Ck/Belgium was determined in chicken embryos as previously described (31). Briefly, embryonated chicken eggs incubated for 10 days (Embryonic Day 10; National Avian Research Facility, The Roslin Institute) were inoculated via the allantoic cavity route with 100 pfu of virus diluted in 100μl of serum-free medium. At 2 days post-infection, embryos were killed by chilling and membrane disruption or membrane disruption and decapitation. Decapitated embryos were washed twice in PBS and fixed for several days in 10% neutral buffered formalin. Five embryos per virus were mounted onto paraffin wax. Embryo tissues were sectioned onto slides and stained with hematoxylin and eosin (H&E) by the Easter Bush Pathology Service. Further unstained sections were used to determine the presence of viral NP by immunohistofluorescence. Deparaffinisation and rehydration, followed by heat-induced antigen retrieval using sodium citrate buffer (10 mM sodium citrate, 0.05% Tween 20, pH 6.0) were performed on tissue sections before staining with primary antibody overnight at 4°C, followed by 3 washes in PBS the next day. Tissue sections were incubated with species-specific secondary antibody and Hoechst for an hour at room temperature. After 3 PBS washes, sections were mounted using ProLong Gold antifade reagent (Invitrogen), scanned using a NanoZoomer XR instrument (Hamamatsu) and analyzed using NDP view, version 2.3, software (Hamamatsu).

### Generation and infection of Chicken 3D intestinal organoids

Embryonic Day 18 Hy-Line Brown Layer eggs were sourced from the National Avian Research Facility, University of Edinburgh, UK. Ethical approvals were obtained from The Roslin Institute’s and University of Edinburgh’s Animal Welfare Ethics Review Board. The humane culling of embryos was conducted under the authorisation of a UK Home Office Project License (PE263A4FA) and adhered to the guidelines and regulations of the UK Home Office Animals (Scientific Procedures) Act 1986.

Duodenum, jejunum, and ileum were extracted from chicken embryos and placed in PBS (Mg^2+^ and Ca^2+^ free) until further processing. For each independent culture, intestines from 3-4 embryos were pooled. Intestinal villi were isolated as described (32). In brief, intestinal tissues were longitudinally opened and cut into 3 mm pieces followed by enzymatic digestion (Clostridium histolyticum Type I Collagenase, 0.2 mg/ml Merck) for 50 minutes at 37°C, 200 rpm. Villi were collected using a 70 µm cell strainer (Corning) by rinsing the inverted strainer followed by centrifugation at 107 x g for 4 minutes. Organoids were seeded in Petri dishes with 12 ml of Floating Organoid Media (FOM), consisting of Advanced DMEM/F12 supplemented with 1 x B27 Plus, 10 mM HEPES, 2 mM L-Glutamine, and 50 U/ml Penicillin/Streptomycin. Organoids were cultured at 37°C with 5% CO_2_.

On day 3 of culture, 200 organoids were seeded in 24-well plates (Corning) in 400 µl of FOM supplemented with or without 50 ng/ml of chicken PLG. Organoids were mock infected or infected with 1000 pfu of viruses according to experimental design, with five wells infected per timepoint. At 1, 24 and 48 hours post-infection, organoids were collected by centrifugation at 10,000 *g* for 2 minutes, and lysed using QIAGEN RLT buffer supplemented with 10 µg/ml β-2-mercaptoethanol (Sigma-Aldrich) and stored at –20°C until use. Cell culture supernatants were collected, frozen on dry ice and stored at –80 °C until use.

### RNA Isolation, reverse transcription and qPCR

RNA was isolated using QAIGEN mini-RNA kits according to the manufacturer’s instructions. Reverse transcription was performed using the Superscript III First Strand Synthesis System (18080051, Thermo Fisher Scientific) according to manufacturer’s instructions using random hexamers and 100 ng of total RNA. The cDNA samples were stored at −20 °C until use. To measure mRNA levels, 1:5 dilution of cDNA was mixed with 10 µl of ABI TaqMan Gene Expression Master Mix (4369016, Applied Biosystems), 1 µl of 20X EvaGreen (31000, Biotum, VWR-Bie & Berntsen) and specific primer pairs for AIV segment 7 and chicken GAPDH (forward and reverse) at a final concentration of 1.15 μM. Reactions were performed in triplicate and no template controls without cDNA were included to detect any potential non-specific amplification. The efficiency of each reaction was calculated based on serial ten-fold dilutions of a calibration sample. The qPCR reaction was carried out at 50 °C for 2 minutes, 95 °C for 10 minutes followed by 40 cycles at 95 °C for 15 s and 60 °C for 1 minute. Primers for AIV segment 7; Forward: CTTCTAACCGAGGTCGAAACGTA, Reverse: CACTGGGCACGGTGAGC (33), and GAPDH; Forward: GAAGGCTGGGGCTCATCTG, Reverse: CAGTTGGTGGTGCACGATG, (Accession number AF047874).

### Sequence analyses

All N1 NA amino acid sequences from viruses of HA subtype H2-H13 and all non-human H1N1 were downloaded from GISAID (34). Human H1N1 sequences were excluded to make the analysis more computationally tractable. These 46,199 sequences were aligned using MAFFT v7.511 using default settings (35, 36). With the resulting alignment, apparent amino acid insertions present in fewer than 5% of sequences were trimmed. Sequences in the resulting trimmed alignment with amino acid coverage below 90% were excluded, leaving 45,001 sequences. The amino acid triplets at positions 130-132 (WSN numbering) were identified. Sequences without complete amino acid information at these positions (*n* = 13) were excluded. Retained sequences were classified as possessing an N-glycosylation motif, Asn-X-Ser/Thr (N-X-S/T) where X is any amino acid except proline (Pro/P), at positions 130-132, or not. Sequences lacking an N-glycosylation motif and without an amino acid other than lysine (Lys/K) at the C-terminal position 453 were defined as potential PLG binders. To place natural sequences possessing this proposed PLG-binding site in the context of the wider diversity of N1 NA proteins, a clustering analysis was used. Within each HA subtype, pairwise amino acid distances were computed from aligned protein sequences and clusters were identified using hierarchical clustering with the nearest chain algorithm (37). Among all sequences lacking the PLG-binding motif, one sequence was randomly selected from each cluster with at least five representatives. This resulted in 119 sequences that were combined with 74 natural sequences possessing a PLG-binding motif for phylogenetic reconstruction. Phylogeny was reconstructed from aligned protein sequences using RAxML-NG (38) with rates of amino acid substitution estimated using the FLU model (39). The best-scoring maximum likelihood tree was mid-point rooted and plotted in R using the ggtree (40).

### Statistical analyses

All graphs were plotted, and all statistical analyses were performed using GraphPad Prism version 10. Individual tests used depended on the data under analysis and are given in figure legends.

## Results

### The unusual Ck/Belgium HA cleavage site reduces susceptibility to trypsin cleavage

To understand the molecular factors contributing to the increased pathogenicity of Ck/Belgium, the HA sequence of Ck/Belgium was compared with the consensus HA sequence of all avian H3 viruses submitted to NCBI from January 2005 to December 2024. This highlighted that Ck/Belgium has a lysine (Figure 1A; highlighted in green) just before the cleavage site instead of an arginine commonly found in H3 viruses. First, we determined if this unusual sequence motif conferred a pseudo HPAIV-like ability of Ck/Belgium to replicate in cells in the absence of trypsin. Chicken lung epithelial CLEC213 cells were infected at low MOI with Ck/Belgium and incubated in serum-free medium with or without trypsin. Subsequent plaque titration in MDCK cells (in the presence of trypsin) showed that Ck/Belgium only replicated in the presence of exogenous protease (Figure 1B). To further understand the effect of the unusual HA cleavage site, we created a Ck/Belgium HA K345R mutant which had the H3 consensus sequence restored and then directly assayed HA cleavage by infecting cells with wild type (WT) and mutant virus followed by western blotting for HA. This showed that both WT and HA K345R needed trypsin for cleavage in CLEC213 cells, as only uncleaved HA0 precursor was visible in the absence of trypsin, while a mix of HA0 and cleaved HA1 and HA2 were present with trypsin (Figure 1C). To provide a more quantitative measurement of trypsin susceptibility, cells were infected with both viruses, then treated with a range of trypsin concentrations and assessed for HA cleavage. From the western blot (Figure 1D), HA K345R showed higher susceptibility to trypsin cleavage than the WT. To examine this in the absence of other viral proteins, cells were transfected with the plasmids encoding WT or mutant HAs and subjected to the same trypsin titration and western blot analysis. Again, the HA K345R mutant showed greater trypsin cleavability (Figure S1). Densitometric measurement of the HA1/HA0 and HA2/HA0 ratios from infection and transfection data confirmed the higher sensitivity of the HA mutant to trypsin (Figures 1E, F), although this was only statistically significant for HA1/HA0 ratios. Thus, the unusual HA sequence of Ck/Belgium does affect trypsin susceptibility, but by reducing cleavability rather than enhancing it.

### The unusual NA and HA sequence polymorphisms in Ck/Belgium affect PLG cleavage of HA

A previous study using another virus isolate from the same Belgium outbreak (A/chicken/Belgium/1940/2019) identified the loss of a glycosylation site at amino acid position 122 on NA (position 130 in WSN numbering) which, by analogy with WSN would allow NA binding of PLG and subsequent trypsin-independent spread (21). Our Ck/Belgium isolate shares the same potential PLG-binding motif in its NA, so we tested whether it too exhibited trypsin-independent spread in the presence of FBS containing PLG. CLEC213 cells were infected with Ck/Belgium, WSN (as a positive control) and Udorn (as a negative control) at low MOI and incubated in the absence or presence of FBS for 24 or 72 hours before staining the cultures with anti-HA to identify infected cells. Individual infected cells were observed for all viruses at 24 hours, with or without FBS (Figure 2A). This pattern of single infected cells remained at 72 hours for all viruses without serum and Udorn with serum. However, clear microplaques were visible at 72 hours for both WSN and Ck/Belgium in the presence of FBS indicating virus spread, thus corroborating the findings of Schon and colleagues (21).

**Figure 2.**
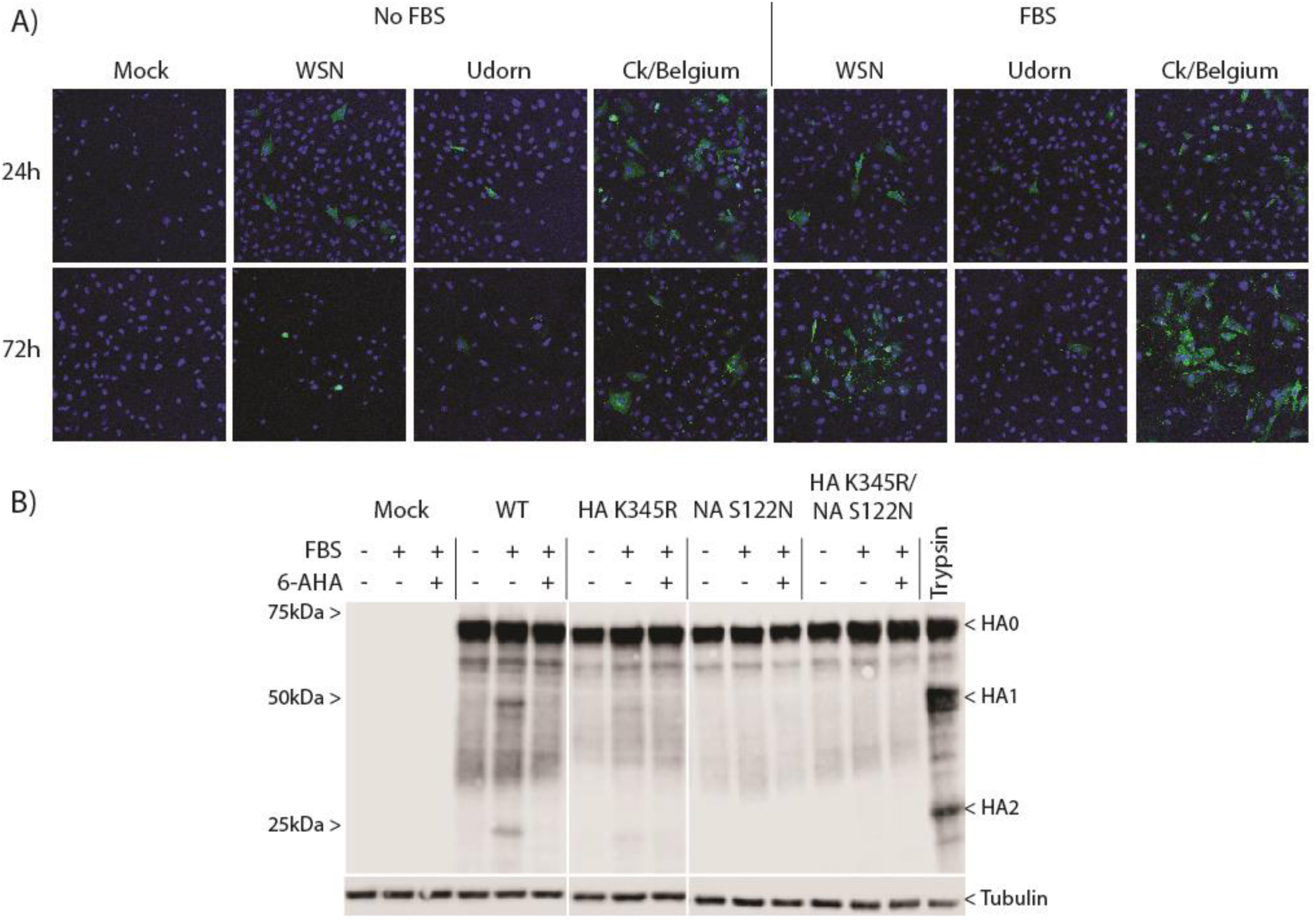
Effects of serum on spread and HA cleavage of Ck/Belgium. (A) CLEC213 cells were infected with WSN, Udorn or Ck/Belgium at an MOI of 0.001 and incubated in the absence or presence of 10% FBS. Cells were fixed and stained with anti-H3 (Udorn and Ck/Belgium) or anti-H1 (WSN) (both in green) and Hoechst (blue) and visualised by fluorescent microscopy. Images for 24 hours and 72 hours post-infection were taken with a 20x magnification lens. (B) CLEC213 cells were infected with WT or mutant viruses at MOI of 3 and incubated in the absence or presence of 10% FBS and 500 μg/ml 6-AHA as indicated. A positive control well was incubated with 1 ug/ml trypsin to provide markers for HA1 and HA2. Cell lysates were harvested 16 hours post-infection and examined by SDS-PAGE and western blotting for HA and tubulin. Positions of molecular mass markers are shown on the left. Images are representative of 2-3 independent experiments.

In order to test the hypothesis that the identity of the amino acid at position 122 on Ck/Belgium NA confers the ability of the virus to spread in the absence of trypsin, as well as to test for any phenotypic association with the unusual HA cleavage site, we created two additional mutants: an NA S122N mutant (restoring the potential glycosylation site) and an HA/NA double mutant, HA K345R/NA S122N. We then infected CLEC213 cells with WT or mutant viruses at high MOI in the absence or presence of FBS and a plasmin-specific inhibitor, 6-AHA, before examining HA cleavage by western blot of cell lysates. Low levels of HA cleavage were observed with the WT virus in the presence of serum, but not in the absence of FBS, nor with FBS further treated with plasmin inhibitor (Figure 2B). The HA K345R mutant showed reduced HA cleavage in the presence of FBS than the WT virus, but was still inhibited by 6-AHA, while the NA S122N and HA K345R/NA S122N viruses did not show detectable HA cleavage under any conditions. These results support the hypothesis that the Ck/Belgium NA recruits PLG for HA cleavage and that the unusual HA cleavage site contributes to the efficiency of this process.

To confirm that HA cleavage resulted from the presence of PLG and not another 6-AHA-sensitive factor present in FBS, we next tested the effects of adding host species-matched purified chicken PLG during virus infections. CLEC213 cells were infected with WT or mutant viruses at high MOI and treated with a range of chicken PLG concentrations, then HA cleavage was determined at 16 hours by western blotting as before. Chicken PLG caused dose-dependent HA cleavage for WT and HA K345R, whereas the HA1/HA2 cleavage products of the NA S122N mutant virus were only faintly observed with 1 μg/ml of PLG (the highest concentration used; Figure 3A). No HA cleavage was detected for the double mutant HA K345R/NA S122N. Quantification of HA1/HA0 (Figure 3B) and HA2/HA0 (Figure 3C) ratios from replicate experiments showed that the HA K345R mutation significantly reduced cleavage compared to the WT HA, as well as confirming only residual cleavage with the NA mutant and no detectable cleavage for the double mutant. To further understand the effect of PLG on virus replication, CLEC213 cells were infected at low MOI and incubated with a range of PLG concentrations. After 24 hours, the number of infected cells were measured by flow cytometry after fixing and staining for surface H3, while the amount of infectious virus released was measured by plaque titration. Spread of the WT virus showed a dose-dependent response to PLG concentration and was significantly higher than all the mutants, particularly viruses with the NA mutation (Figure 3D). Titres of infectious virus also increased with PLG concentration and again, all mutations reduced virus replication (Figure 3E). These results indicate that the unusual Ck/Belgium HA cleavage site and NA recruitment of PLG collaborate to induce HA cleavage and virus spread *in vitro*.

**Figure 3.**
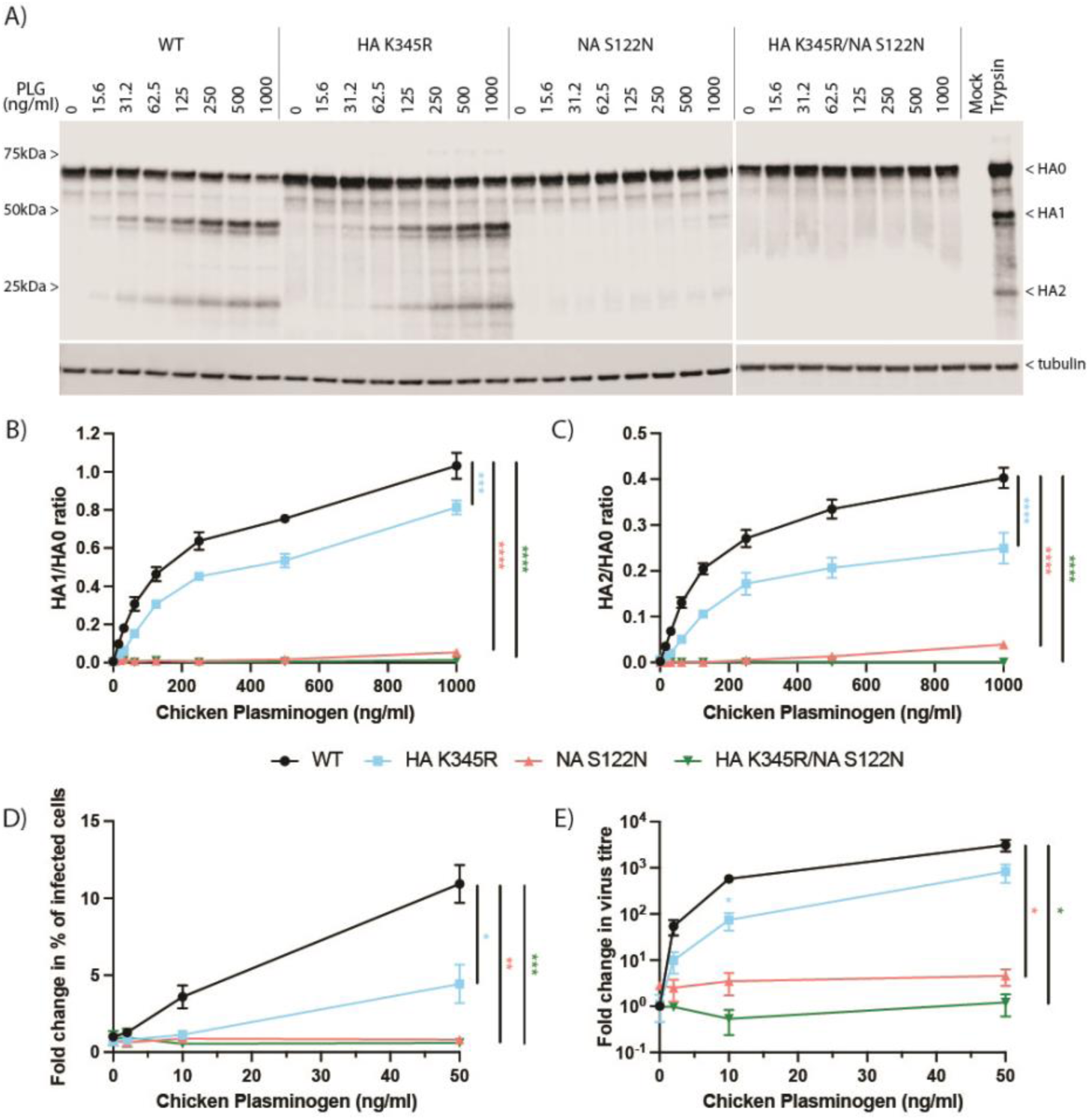
Synergy between the Ck/Belgium HA cleavage site and NA PLG recruitment motif. (A-C) CLEC213 cells were infected with WT or mutant viruses at an MOI of 3 and incubated in the presence of a range of chicken PLG concentrations for 16 hours. (A) Cell lysates were examined by SDS-PAGE and western blotting for HA and tubulin. Positions of molecular mass markers are shown on the left. (B, C) Ratios of cleaved to uncleaved HA products from replicate blots were quantified by densitometry. Data are mean ± SEM of 3 independent experiments. (D, E) CLEC213 cells were infected with WT or mutant viruses at an MOI of 0.001 and incubated with the indicated amounts of PLG for 24 hours. (D) Cells were fixed and stained with anti-H3 and the number of infected cells quantified by flow cytometry, while (E) released infectious virus was titred by plaque assay. Data are presented as fold increase induced by PLG relative to WT virus without PLG and values are mean ± SEM of 3-4 independent experiments. Statistical analyses are area under the curve for each virus (B-E) or one-way ANOVA and Dunnett’s multiple comparisons test for HA K345R in (E).

### Synergy between the HA cleavage site and NA-mediated PLG recruitment is not cell type-dependent and occurs *ex vivo* and *in ovo*

PLG activation is a tightly regulated process with multiple cell-surface stimulatory and inhibitory factors, whose abundance can vary by tissue or cell type (41). All our experiments so far were performed in CLEC213 cells, so it was therefore important to test the significance of the Ck/Belgium HA and NA sequence motifs in other avian cells. Using a single concentration of 250 ng/ml PLG, cleavage of WT Ck/Belgium HA was again significantly higher than HA K345R and no cleavage was found for NA S122N and HA K345R/NA S122N in infected CLEC213 cells (Figure 4A). Very similar results were obtained using another chicken cell line (DF-1 fibroblasts; Figure 4B), as well as in duck (CCL-141; Figure 4C) and quail (QT-35; Figure 4D) fibroblast cell lines. Thus, the HA processing induced by the collaborative effects of the unusual HA cleavage site and NA-recruited PLG were not specific to a single type of chicken cell but also applied to immortalised cells from other avian species.

**Figure 4.**
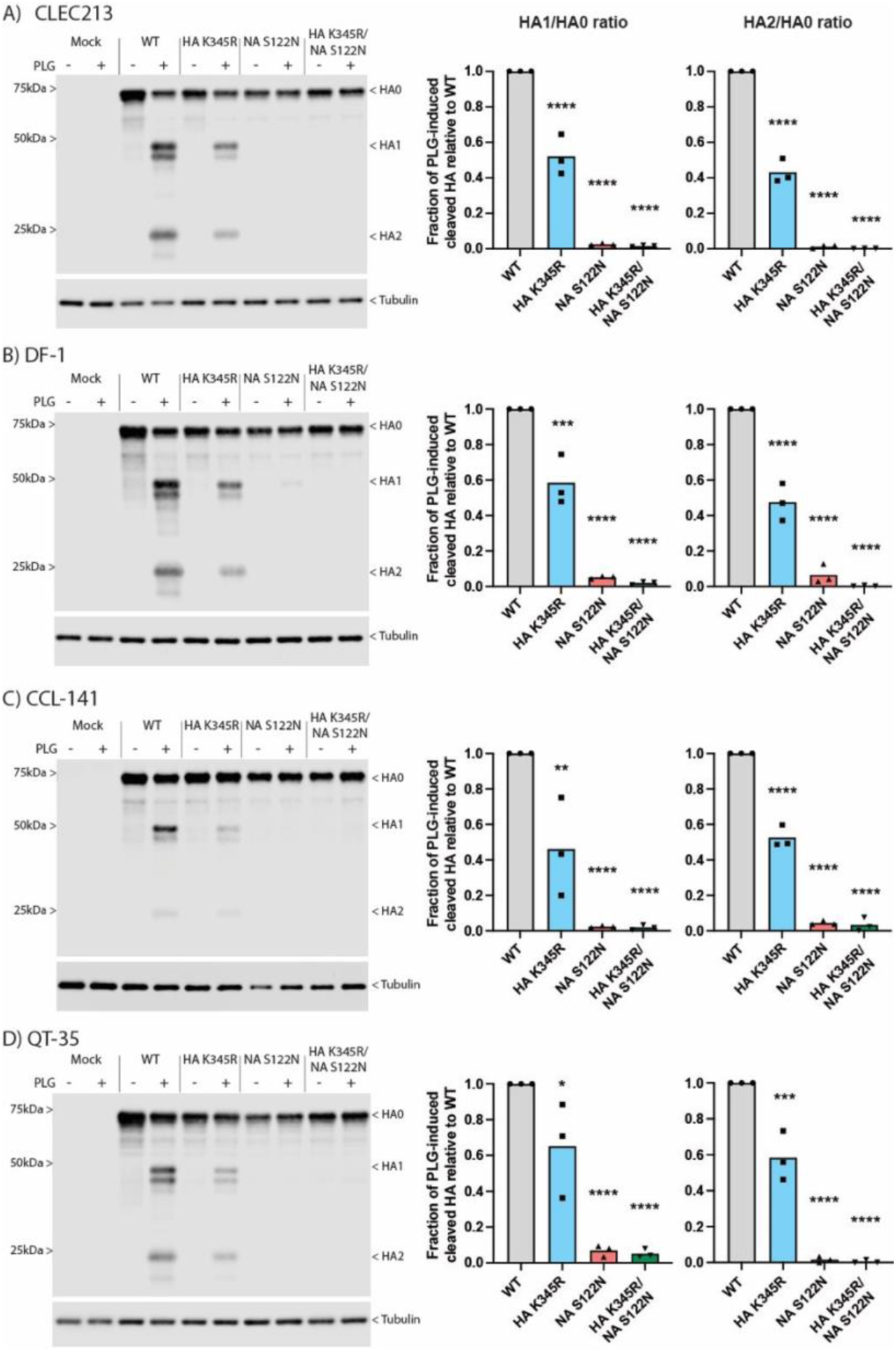
The effects of Ck/Belgium HA cleavage site and NA PLG-binding motifs are not cell type or species-specific. PLG-induced HA cleavage were determined in chicken (A, B), duck (C) and quail (D) cell lines. Cells were infected with WT or mutant viruses at MOI of 3 for 16 hours in the absence or presence of 250 ng/ml PLG before analysis by western blot for HA and tubulin. Images are representative of 3 independent experiments. Dots on graphs represent densitometric ratios of HA1/HA0 and HA2/HA0 from individual experiments, normalised to the respective WT values. Bars indicate the means of 3 independent experiments. One-way ANOVA followed by Dunnett’s multiple comparisons test were performed for statistical analyses. * = p < 0.05, ** = p < 0.01, *** = p < 0.001, **** = p < 0.0001.

In order to further examine the biological relevance of PLG-mediated HA cleavage, we used chicken organoids as a complex cell model of infection in primary cells (32). Groups of approximately 200 organoids pooled from 3-4 chick embryos were infected with WT or mutant viruses in the absence or presence of PLG. Virus replication was examined 24 hours later by quantifying the intracellular expression of segment 7 RNA as well as virus released into the supernatant. The addition of PLG increased segment 7 accumulation significantly in WT and HA K345R infections whilst the NA S122N mutation suppressed this PLG-induced increase, both alone and with the HA mutation (Figure 5A). Although variable between batches of organoids, the addition of PLG also significantly increased the titre of released WT, but not mutant viruses (Figure 5B). Overall, these data further corroborate the importance of the HA cleavage site and NA PLG-binding motifs for Ck/Belgium replication.

**Figure 5.**
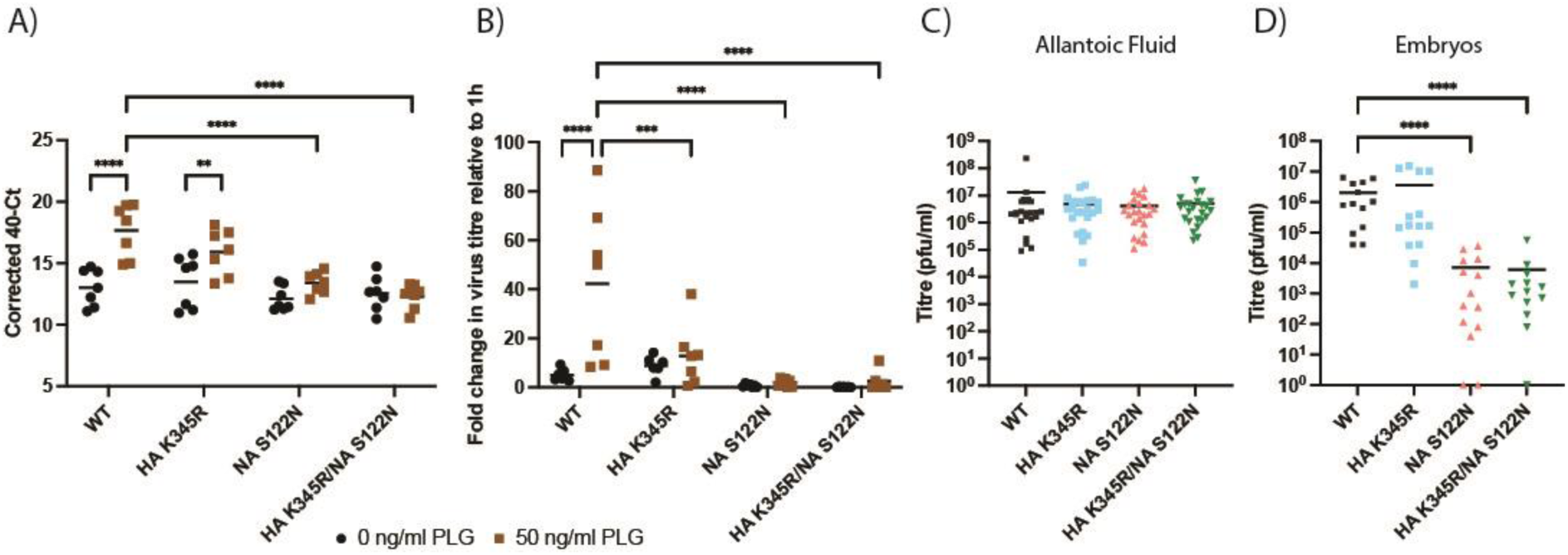
The effects of Ck/Belgium HA cleavage site and NA PLG-binding motifs in *ex vivo* and *in ovo* systems. (A, B) Chicken organoids were infected with 1000 pfu of WT or mutant viruses in the absence or presence of 50 ng/ml PLG for 24 hours. Virus replication was measured by (A) qPCR of intracellular segment 7 or (B) plaque assay of released virus. Each dot represents a pool of organoids derived from 3-4 chicken embryos and lines represent the means of 6-7 organoid pools from 2 independent experiments. Two-way ANOVA (qPCR) or mixed-effects analysis (plaque assay) followed by Dunnett’s multiple comparison test were performed for statistical analyses. (C, D) The allantoic cavities of 10-day old embryonated chicken eggs were inoculated with 100 pfu of WT or mutant viruses and incubated for 2 days. Virus titres in (C) allantoic fluid and (D) chicken embryos were examined. Dots represent virus titres from at least 13 individuals and lines indicate the means from 2 independent experiments. Statistical differences were determined using Kruskal-Wallis test followed by Dunn’s multiple comparisons test.

As a further *in ovo* model of infection, we inoculated the allantoic cavity of 10-day old embryonated chicken eggs with WT and mutant Ck/Belgium viruses. There were no differences in the titres of the various viruses harvested from the allantoic fluid 48 hours later (Figure 5C). WT Ck/Belgium and the HA K345R mutant also replicated well in the chicken embryos, but in contrast, NA S122N and HA K345R/NA S122N showed notably lower virus titres (Figure 5D). These results were also supported by immunostaining of the chicken embryos for virus antigen. In lung, heart and brain sections, NP expression was only observed for WT and HA K345R viruses, whereas the staining for NA S122N and HA K345R/NA S122N was similar to that of mock-infected embryos (Figure 6). These findings indicated the importance of NA-mediated PLG recruitment in the pathogenesis of Ck/Belgium.

**Figure 6.**
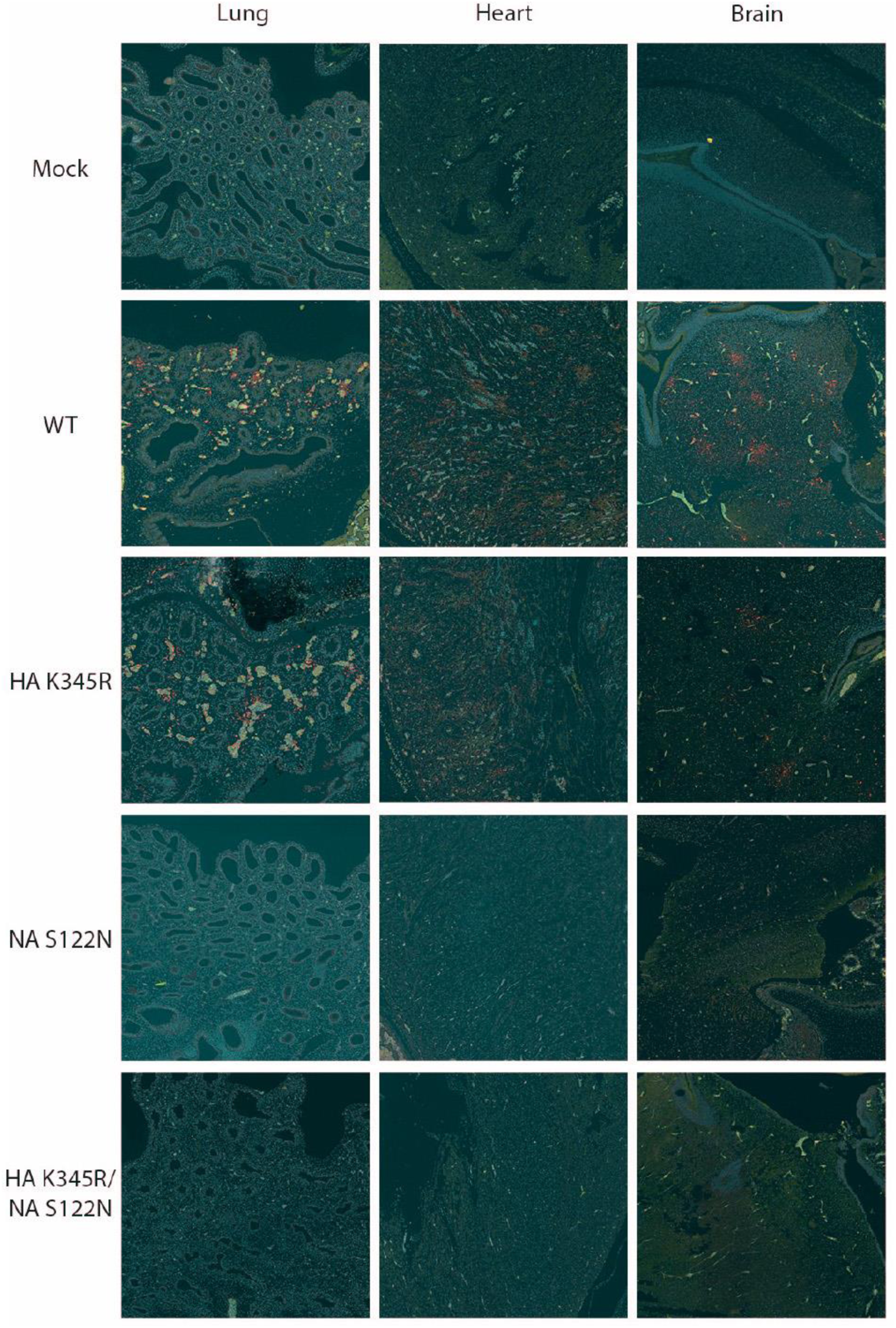
The Ck/Belgium NA PLG-binding motif facilitates systemic replication in the chick embryo model. The allantoic cavities of 10-day old embryonated chicken eggs were inoculated with 100 pfu of WT or mutant viruses and incubated for 2 days. Embryos were fixed, sections cut and then immunofluorescently stained for viral NP and cellular nuclei using Hoechst dye. Images show the lung, heart and brain of chicken embryos infected with the indicated viruses are representative of 10 embryos per condition. Red – NP, blue – nucleus, green – autofluorescence.

### PLG recruitment by N1 NA has also evolved in moderately virulent H6 subtype LPAIVs

Schon and colleagues previously searched the Influenza Research Database for other N1 NA sequences that contained a PLG-binding motif, identifying 15 independent virus isolates not linked to the Belgian 2019 H3N1 outbreak (21). Of these, the largest category (discounting multiple sequences of the lab-adapted WSN, PR8 and a reassorted derivative of the latter) were human infections, while only two were independent isolates of LPAIVs, both from the 1980s. To further investigate the emergence of the PLG-binding motif across N1 NAs, we carried out a similar search using the Global Initiative on Sharing All Influenza Data (GISAID) database (34). From over 38,000 sequences that met inclusion criteria (we excluded human seasonal viruses for computational tractability), we found 165 sequences lacking the NA glycosylation site and with a C-terminal lysine which may permit PLG binding. Of these, 153 were inferred to be from individual naturally occurring virus isolates (Tables 1 and S1). These potential PLG-utilising viruses span a date range from 1983 to 2025, possess H1, H3, H5 or H6 subtype HAs and come from a range of host species, both mammalian and avian, but predominantly domestic poultry (Table 1). To gain an understanding of the distribution of these PLG-binding viruses among the wider diversity of N1 NAs, they were placed in a phylogeny representative of protein diversity. Figure 7A highlights NAs with PLG-binding motifs as well as indicating subtype and the associated host types across the tree. Most (100 or 83.3%) were H3N1 sequences with the vast majority belonging to the 2019 Belgium outbreak (*n* = 99, of which 20 are plotted in Figure 7A) and one other sequence (A/duck/Buryatia/664/1998; identified in (21)) being highly diverged from this outbreak. Outside of H3N1 strains, the next largest group of NAs possessing the PLG-binding motif were H6N1 viruses (31 sequences). We noted two clusters of H6N1 viruses in which the motif indicative of PLG-binding appears to have emerged independently. Based on the dates and countries which the isolates were collected, they derive from two epizootic clusters: 3 sequences representing a 2010 outbreak in the Netherlands (4) while the other 28 sequences are from linked but genetically distinct outbreaks in Ireland, Northern Ireland and Scotland (5). All four outbreaks were associated with notable drops in egg production and increased mortality. For the Ireland/UK outbreak, there was also evidence of systemic virus spread within the birds (5). We also found H5N1 and H1N1 subtype viruses (15 and 7 sequences, respectively) that possessed the PLG-binding motif. These tend to occur as singletons in the tree, suggesting independent evolution, except for two clusters of viruses sampled as part of the ongoing epizootic of HPAI H5N1 in the USA (Table 1). However, closer examination of genotypes and NA sequences involved suggested that these also represented several independent emergence events. The majority of other H5N1 strains with potential PLG-recruiting NAs are also HPAIVs. Thus, the N1 PLG-binding motif has evolved independently on multiple occasions, but most notably in LPAIV outbreaks in poultry, in the context of virus strains exhibiting increased pathogenicity.

**Figure 7.**
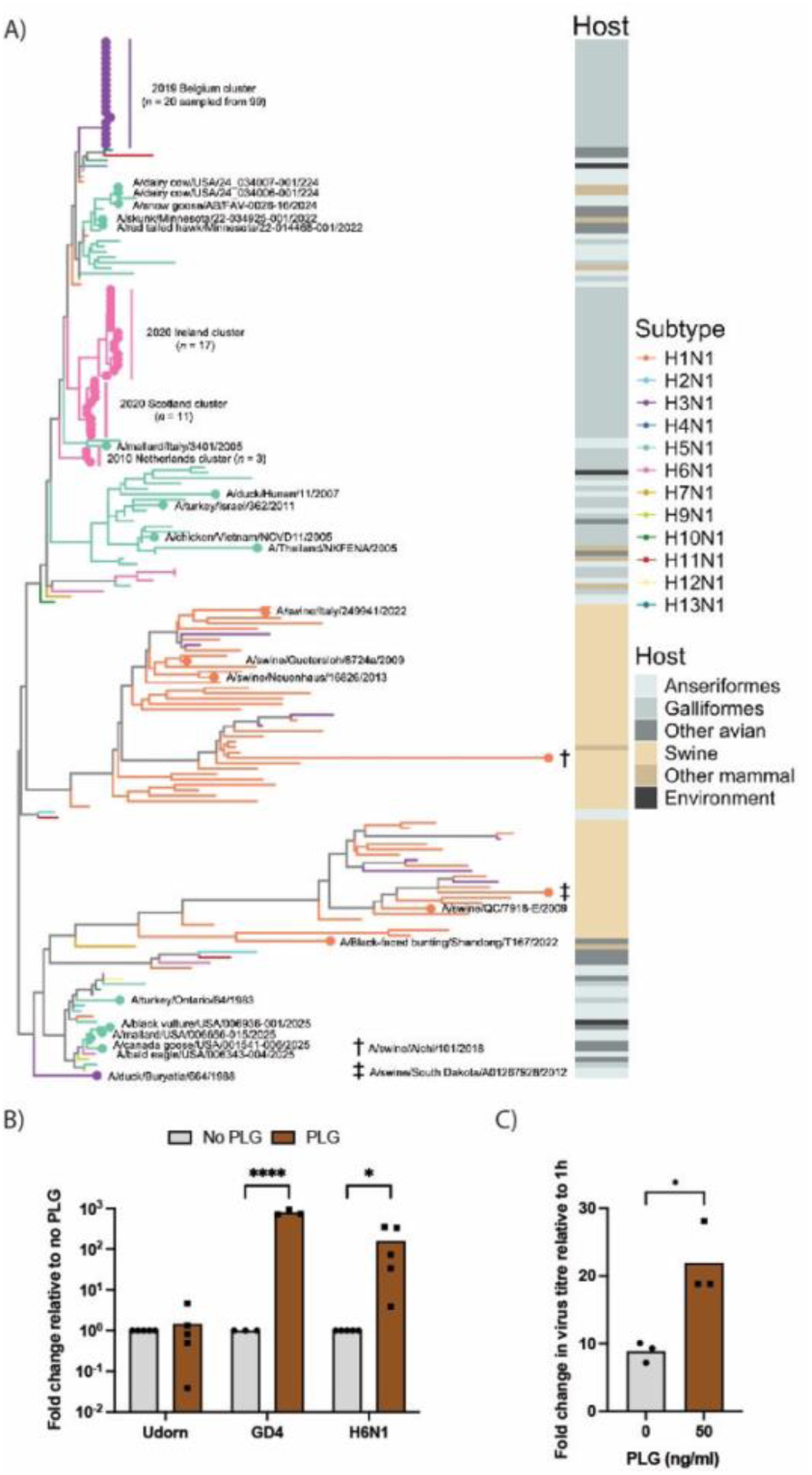
Identification of potential NA-mediated PLG recruitment in other N1 subtype viruses. (A) NA phylogeny reconstructed from amino acid sequences of 74 N1 NAs with a PLG-binding motif and 119 other sequences representing the diversity of N1 NAs. The tree is mid-point rooted and branches are coloured by subtype. The positions of viruses with PLG-binding motifs are highlighted by circles; virus names or epidemiological clusters are labelled. Alongside, a coloured column indicates host type which includes the avian orders Anseriformes (ducks, geese and swans) and Charadriiformes (gulls, waders and auks). (B) Experimental tests of PLG utilisation by a 2010 Netherlands H6N1 outbreak virus. CLEC213 cells were infected with Udorn, Ck/Belgium or H6N1 viruses at MOI=0.001 for 48 hours in the absence or presence of 50 ng/ml PLG. (C) Chicken organoids were infected with H6N1 virus in the absence or presence of 50 ng/ml PLG for 24 hours. Virus replication in (B) and (C) was examined by plaque assay of released virus. Dots represent biological replicates in (B) or pools of organoids from 3-4 chicken embryos in (C) and bars indicate the means. Statistical differences were determined using 2-way ANOVA followed by Sidak’s multiple comparisons test in (B) and unpaired t-test in (C).

**Table 1.**
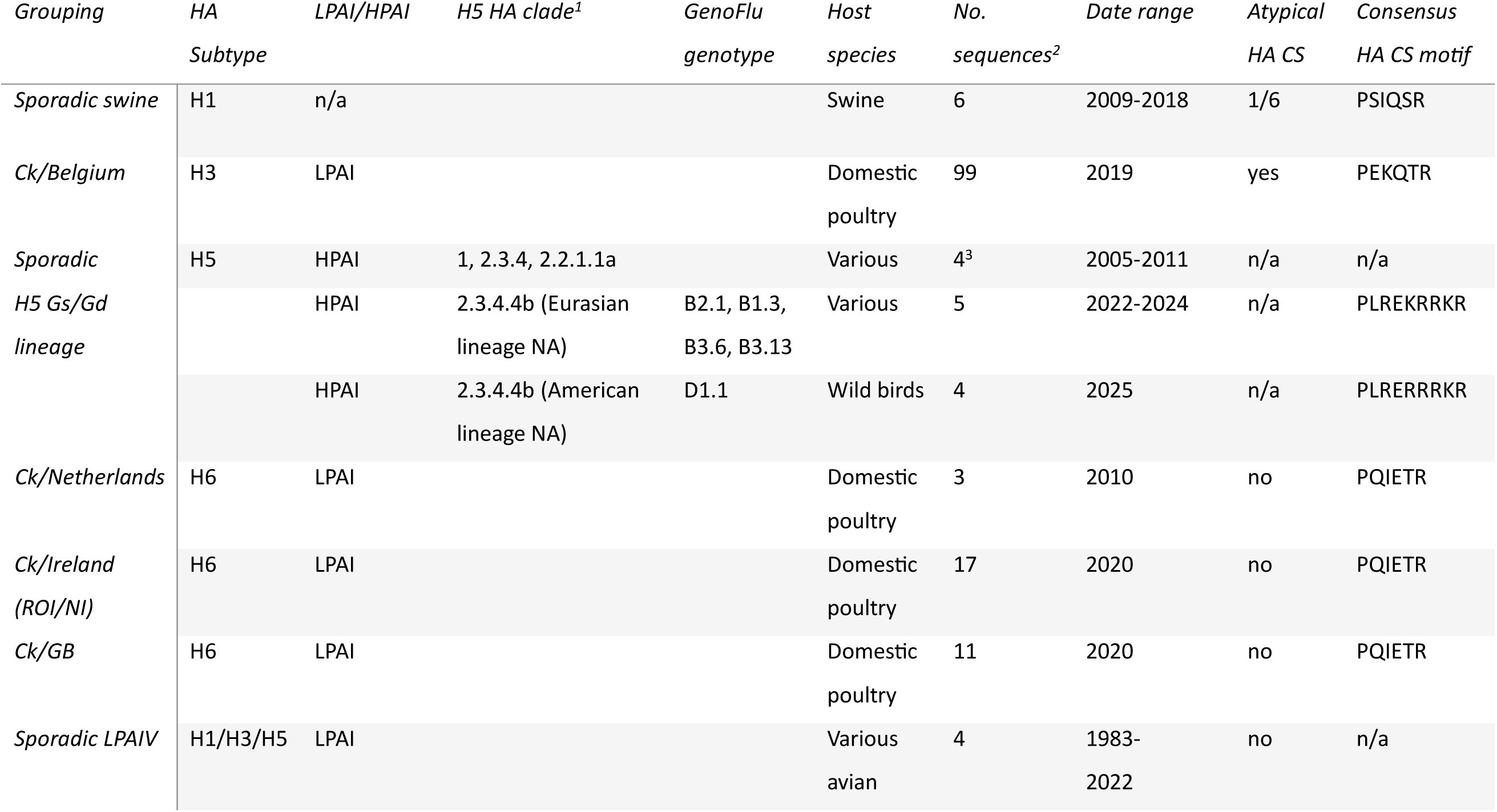
Categories of non-laboratory adapted influenza A virus N1 strains known or suspected to recruit PLG. ^1^For H5 HA clade 2.3.4.4b, viruses are distinguished as possessing either neuraminidase of either Eurasian or American AIV lineages. ^2^Number of natural viruses with PLG-binding motifs identified in analysis of 44,988 N1 sequences (10,695 H1N1, 69 H2N1, 310 H3N1, 36 H4N1, 32,629 H5N1, 727 H6N1, 2 H8N1, 48 H9N1, 90 H10N1, 63 H12N1 and 6 H13N1). ^3^One isolate lacks an associated HA sequence.

We had an isolate of the 2010 Netherlands H6N1 outbreak available (A/chicken/Netherlands/917/2010). Accordingly, to determine if this virus could utilise PLG, CLEC213 cells were infected at low MOI with Udorn, Ck/Belgium or the H6N1 LPAI viruses and incubated without trypsin but in the absence or presence of PLG for 48 hours. Plaque titration of released virus showed that addition of PLG significantly increased the virus titres of Ck/Belgium and H6N1, but not Udorn (Figure 6B). Furthermore, PLG addition also significantly increased replication of the H6N1 virus in chicken organoids (Figure 6C). We conclude that similarly to Ck/Belgium, the atypical N1 sequence of the Netherlands H6N1 outbreak also leads to PLG-mediated spread of the virus, potentially explaining the higher than normal pathogenicity of the outbreak viruses.

## Discussion

Despite being categorised as an LPAI, H3N1 viruses from the Ck/Belgium family have been characterised as moderately virulent in poultry, showing systemic replication as well as causing appreciable mortality and a severe drop in egg production (8). The unexpected pathogenicity of the outbreak strain has been linked to PLG-cleavage of the HA, potentially mediated by loss of a glycosylation site in NA that, by analogy with the laboratory-adapted WSN and PR8 strains, leads to recruitment of PLG to the cell surface by NA (18, 21, 42). Here, we report molecular tests of this hypothesis. We confirmed PLG-driven HA cleavage and virus spread of Ck/Belgium using purified chicken PLG in avian cell systems from three species, including fibroblast, epithelial and organoids. Using reverse genetics, we show that the key determinants of this are indeed the loss of a potential glycosylation site in NA, along with a variant monobasic cleavage site in the H3 HA that potentiates PLG cleavage and increases virus replication *in vitro*, as well as *ex vivo* and in *in ovo* systems. Furthermore, we identify further outbreaks of unexpectedly pathogenic H6N1 LPAIVs with similar traits, suggesting this is a general virulence mechanism for N1 LPAIVs.

This study brings the number of virus lineages experimentally verified to use PLG-recruitment by an N1 NA for HA processing to four: the H1N1 WSN and PR8 families, the 2019 Belgium H3N1 outbreak and the 2010 Netherlands H6N1 LPAIV outbreak. WSN (18, 19, 22) and now the Belgian H3N1 viruses have been shown to depend primarily on the N1 mutation that destroys a potential N-linked glycosylation site, but to also possess a further adaptive mutation in the HA cleavage site that enhances plasmin-mediated proteolysis. Although both sequence motifs are important, the NA change is clearly more consequential, as its reversal to normal consensus has a greater effect on replication *in vitro* and pathogenicity *in ovo* than restoring the HA cleavage site to the more usual sequence ((18, 19, 22) and this study). Indeed, the NA PLG-binding polymorphism was sufficient to correctly identify the Netherlands H6N1 outbreak virus as using PLG, although its HA has the usual H6 PQIETR/GLF cleavage site sequence (Table 1). The viruses from the linked 2020 outbreaks of moderately virulent H6N1 LPAIVs in Irish and UK poultry flocks (4, 5) also have a normal HA consensus cleavage site, but nevertheless we think it is plausible that they too will prove to be able to use PLG to mature HA.

Whether the acquisition of an unusual HA cleavage site sequence represents the “fine-tuning” of the PLG-recruitment virulence mechanism or is selected for other reasons (perhaps HA subtype-specific) remains to be determined. The unusual cleavage site in WSN did not obviously alter its ability to be cleaved by trypsin (22) but the Ck/Belgium HA showed a moderate reduction in trypsin-cleavability. In mammals, PLG is both broadly expressed and secreted from the blood into other tissues, including the lung and central nervous system (43, 44), but it is unclear whether a virus that <u>can</u> use PLG <u>only</u> uses PLG, or (more plausibly perhaps), HA is simply cleaved by whatever protease with the right specificity comes along first. If the latter is true, then it would be interesting to test whether the Ck/Belgium HA cut site also affects cleavage by airway proteases and thus represents a balancing act between transmissibility and systemic replication.

Further relating to the diversity of proteases involved in HA cleavage is the intriguing number of H5 HPAIV HAs, primarily from the A/Goose/Guandong/1/1996 lineage, which have an N1 NA without the key glycosylation site (Table 1). At first sight, it seems unlikely that a virus that has evolved a highly efficient furin-cleavable HA would gain an advantage from being able to also use PLG. Furin-like proteases that cleave HA are usually viewed as ubiquitous in cell types and tissues, explaining the systemic replication of H5 and H7 HPAIVs in poultry (2, 16, 45). However, furin-like proprotein convertase subtilisin-kexin (PC/PCSK) genes are a large family that vary with respect to secretion versus transmembrane residence and intracellular localisation, as well as ability to cleave HA (46). It is therefore possible that there are circumstances in which PLG recruitment might be advantageous even in an HPAIV background. Given that PLG recruitment is well correlated with neurovirulence (18) and that several of the recent HPAIV isolates with the potential to recruit PLG are Clade 2.3.4.4b viruses isolated from mammals, including cows (Table 1), determining whether this loss of glycosylation in some HPAIV N1 NAs is genuinely functional or mere sequence noise could be important for understanding the evolution of pathogenicity of this newly emergent variant.

Overall, our study reveals that the unusual HA sequence of Ck/Belgium works in concert with NA-recruited PLG to increase HA cleavage, virus spread, and systemic replication in avian cells. We also provide evidence from another outbreak-associated LPAI H6N1 virus that this is potentially a generalisable virulence mechanism for LPAIVs. We suggest that future outbreaks of unexpectedly pathogenic LPAIVs with an N1 NA in poultry should have their NA and HA genes monitored for loss of the glycosylation site and secondarily, an atypical HA cleavage sequence.

## Supporting information

Table S2 – GISAID acknowledgments

## Acknowledgments

We thank Prof Massimo Palmarini (MRC-University of Glasgow Centre for Virus Research, UK) for MDCK/chicken ANP32A cells, Dr Laurence Tiley (University of Cambridge, UK) for QT-35 cells, Dr Sascha Trapp (French National Institute for Agriculture, Food and Environment, France) for CLEC213 cells, Dr Leah Goulding (University of Nottingham, UK) for CCL-141 cells, Prof Janet Daly (University of Nottingham, UK) for anti-H3 antibody and Prof Richard Compans (Emory University, Atlanta) for Udorn virus. We are grateful to the staff of the Roslin Institute’s Central Services Unit, Facilities team, Bioimaging facilities, Easter Bush Pathology and to the National Avian Research Facility for supply of eggs. We also gratefully acknowledge all GISAID data contributors, i.e., the Authors and their Originating laboratories responsible for obtaining the specimens, and their Submitting laboratories for generating the genetic sequence and metadata and sharing via the GISAID Initiative, on which this research is partially based.

This work was funded by Institute Strategic Program Grant BBS/E/RL/230002C and BBS/E/RL/230002D from the Biotechnology and Biological Sciences Research Council (BBSRC) to PD, LV, SL, RMP and EG, and BB/V019899/1 “FluNuance” from the BBSRC under the aegis of the International Coordination of Research on Infectious Animal Diseases (ICRAD) programme to SdW, PD and LV.

## Supplementary information

Table S1; Lists of N1 NA sequences on the GISAID database lacking a glycosylation site at position 130 (WSN numbering). Table S1 tab lists sequences judged to be from independent natural virus isolates. Sequences from laboratory adapted/manipulated strains are listed in a separate tab.

Table S2; GISAID acknowledgments for specific N1 sequences

Figure S1; Trypsin-dependent cleavage of HA in transfected CLEC213 cells

---

**Figure S1.**
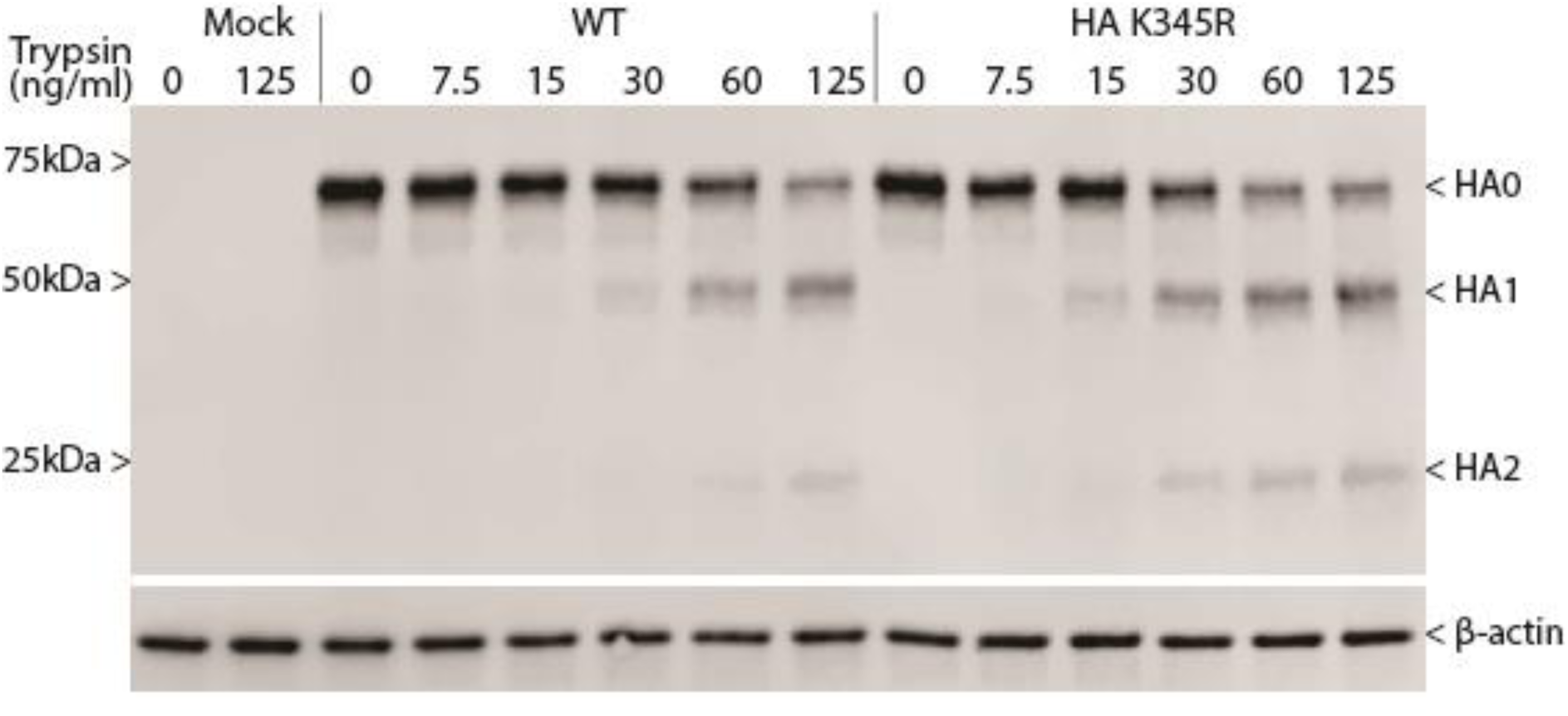
Trypsin-sensitivity of Ck/Belgium HA in transfected cells. CLEC213 cells were transfected with plasmids encoding WT or K345R HA for 48 hours and the indicated trypsin concentrations were added for the last 2 hours. SDS-PAGE and western blotting were performed to detect HA and ß-actin. Positions of molecular mass markers are shown on the left.

